# Mental Imagery abilities affect visual working memory performance: evidence from aphantasic participants

**DOI:** 10.1101/2025.08.08.669378

**Authors:** Chong Zhao, Edward K. Vogel, Wilma A. Bainbridge

## Abstract

Visual imagery refers to the mental generation of visual representations of stimuli, while visual working memory involves retaining visual information for a short period without external input. Due to the conceptual overlap between these two constructs, successful performance on visual working memory tasks may rely on the use of visual imagery to rehearse items during the retention interval. Consequently, individuals with aphantasia, who lack voluntary visual imagery, may experience difficulties with such tasks.

However, prior research has suggested that some individuals with aphantasia might employ non-visual strategies to compensate for this deficit. In two experiments, we examined visual working memory performance in aphantasic and control participants across a range of stimulus types. In Experiment 1, participants completed a change localization task using color squares and complex fractals; in Experiment 2, stimuli included real words, phonologically valid pseudowords, and phonologically invalid pseudowords. Across both experiments, aphantasic participants demonstrated significantly impaired visual working memory compared to controls. Notably, their performance was equally impaired for stimuli that were easily verbalizable (i.e., colors and words) and those that were not (i.e., fractals and pseudowords). Furthermore, individual differences in visual imagery ability, as measured by the Vividness of Visual Imagery Questionnaire (VVIQ), significantly predicted working memory performance across all stimulus types. These findings provide direct evidence for the critical role of visual imagery in supporting visual working memory.

## Introduction

Mental imagery refers to the capacity to generate and manipulate visual representations of objects or scenes in the absence of direct sensory input. A closely related construct, visual working memory, involves the temporary maintenance of visual information over brief periods during which the stimuli are no longer visible. These two constructs not only share conceptual similarities (Tong, 2013) but are also supported by overlapping neural substrates. For instance, neuroimaging evidence suggests that visual working memory and visual imagery are represented similarly in early visual areas (Albers et al., 2013).

Recent research on mental imagery has identified a subset of individuals who report an inability to voluntarily generate visual mental imagery, a condition now referred to as aphantasia. Despite having intact vision and no apparent deficits in recognition, individuals with aphantasia are unable to form visual images in their "mind’s eye." Prevalence estimates suggest that aphantasia affects approximately 2–5% of the population, highlighting its relevance as a relatively common but underexplored cognitive variation (Keogh & Pearson, 2018; Zeman et al., 2015, p. 2).

Despite growing interest in aphantasia, there is inconsistent evidence regarding how this condition affects visual working memory (VWM) performance. One possible hypothesis is that mental imagery abilities play an important role in visual working memory tasks, and thus an impairment in mental imagery would result in an impairment of visual working memory. Supporting this hypothesis, a previous study using a binocular rivalry paradigm found that individuals with stronger mental imagery abilities tended to perform better on visual working memory tasks (Keogh & Pearson, 2011). In the experiment, participants were asked to remember a single grating stimulus over a brief delay and were then presented with a binocular rivalry display as a memory probe. The researchers found that participants with stronger visual imagery were more likely to perceive the remembered image as dominant, indicating an interaction between working memory and sensory-level imagery. Furthermore, the same study demonstrated that when background luminance was reduced, a manipulation known to disrupt visual imagery, individuals with strong imagery abilities showed impaired performance on the working memory task, while those with weaker imagery abilities were unaffected. This suggests that visual working memory performance may partially depend on imagery-based sensory representations. In a separate case study, an individual with aphantasia performed significantly worse than IQ-matched controls on a demanding visual working memory task that asked participants to memorize the boundary of a single presented shape over 4 seconds of delay (Jacobs et al., 2018). Together, these findings suggest that aphantasia may be associated with deficits in visual working memory, particularly under conditions that place high demands on internal visual representations.

However, surprisingly a growing number of other studies have found that individuals with aphantasia perform comparably to controls on visual working memory tasks, despite the absence of visual imagery. This counterintuitive pattern has led researchers to suggest that aphantasics may compensate by relying on non-visual strategies, such as verbal encoding or spatial reasoning, to support task performance. For instance, in the same case study we discussed above, the individual with aphantasia performed similarly to the IQ-matched controls on an easier working memory task, specifically in detecting a big change in visual boundaries between the encoding and test phase (Jacobs et al., 2018). Keogh, Wicken, and Pearson (2021) reported that although aphantasic participants demonstrated similar visual working memory capacity to controls, their strategies were predominantly non-visual based on their self-reported strategies, in contrast to the visual strategies used by control participants. Consistent with this, Knight, Milton, and Zeman (2022) found that aphantasic individuals performed on par with controls on a visual working memory task that asked participants to report the orientation of a Gabor pattern even when a distractor was introduced between encoding and retrieval. Similarly, Pounder et al. (2022) found that aphantasic participants performed comparably to control subjects on three visuo-spatial working memory tasks: the Spatial Span task, the Mental Rotation task, and the One Touch Stockings of Cambridge task. Furthermore, a recent neuroimaging study revealed that an aphantasic participant exhibited neural representations during a visual memory task that were strikingly similar to those of their identical twin, who had intact visual imagery abilities (Megla et al., 2025). Collectively, these findings suggest that while aphantasics may lack visual imagery, they may compensate by engaging alternative, non-visual cognitive strategies to maintain information in working memory.

A recent challenge to the verbal compensation hypothesis for aphantasics arises from the expectation that if individuals with aphantasia rely predominantly on verbal strategies, their performance on verbal memory tasks should be comparable to that of control participants, as these tasks do not depend on visual imagery. However, recent findings suggest otherwise. Aphantasic individuals have been shown to perform worse than controls on both verbal short-term memory tasks (when researchers played audio recordings of word lists) and visual short-term memory tasks (when researchers presented the word lists visually, Monzel et al., 2022).These results complicate the interpretation of strategy use in aphantasia during visual memory tasks.

One challenge in interpreting previous findings on visual working memory (VWM) in aphantasia is the wide variety of stimuli and paradigms that have been used across studies. These differences make it difficult to directly compare results or draw firm conclusions. Recent research has clarified that VWM is composed of distinct components, namely, capacity and precision, which should be measured separately (Zhang & Luck, 2008). Under the assumption of this dual-component model, other research had shown that more complex stimuli, such as polygons and Japanese characters, tend to increase confusion errors during the decision-making stage, rather than reflect true storage limitations (Awh et al., 2007). Mapping onto the two-factor model of working memory, more complex stimuli like Gabor patterns used in prior research with aphantasia could be tapping into precision measures instead of working memory capacity. Given our primary interest in assessing the capacity of VWM in individuals with aphantasia, it is essential to use a task that is both reliable and sensitive to differences across conditions and individuals. To address this need, we employed the change localization task, a paradigm recently shown to robustly measure VWM capacity (Zhao et al., 2022). In this task, participants briefly view an array of visual items, and after a short delay, are asked to identify which single item has changed in the test array. This task has demonstrated high split-half reliability (Spearman-Brown correlation > 0.75) with just 50-60 trials, taking approximately five minutes to complete. Therefore, in the present study, we used this well-validated change localization task with a range of visual stimulus types to assess VWM performance in aphantasic and control participants.

In the present study, we investigated whether visual working memory is impaired in individuals with aphantasia. In Experiment 1, we administered a change localization task with either color squares or fractals as stimuli. The color stimuli were highly distinct and easily nameable, whereas the fractals were more visually complex and difficult to verbally label. This design allowed us to examine whether aphantasic participants, who may rely more heavily on verbal coding strategies, would perform more poorly on the fractal task compared to the color task, relative to control participants. In Experiment 2, we aimed to further explore the potential use of non-visual strategies in compensating for visual working memory deficits in aphantasia. To generalize our findings to visually presented words and letter strings, participants completed a change localization task involving words and two types of pseudowords: phonologically valid and phonologically invalid. We hypothesized that if aphantasic individuals compensate through phonological coding, they would show preserved performance on tasks involving words and phonologically valid pseudowords than on those with phonologically invalid pseudowords compared to control participants.

## Experiment 1

### Methods

#### Participants

We assessed mental imagery ability in both the aphantasia and control groups using the Vividness of Visual Imagery Questionnaire (VVIQ; Marks, 1973), a self-report measure that evaluates the vividness of an individual’s visual mental imagery. VVIQ scores range from 16 to 80, with higher scores indicating more vivid imagery. Following the criteria from Bainbridge et al. (2021), participants scoring 25 or below were classified as having aphantasia. For the control group, only individuals scoring above 40 were included (see **Fig. 1A**). As part of the online task, we included attentional check questions to ensure participant engagement. These consisted of simple classification items such as “Is a coffee mug edible?” with response options: press ‘E’ for edible and ‘I’ for non-edible.

**Fig 1.**
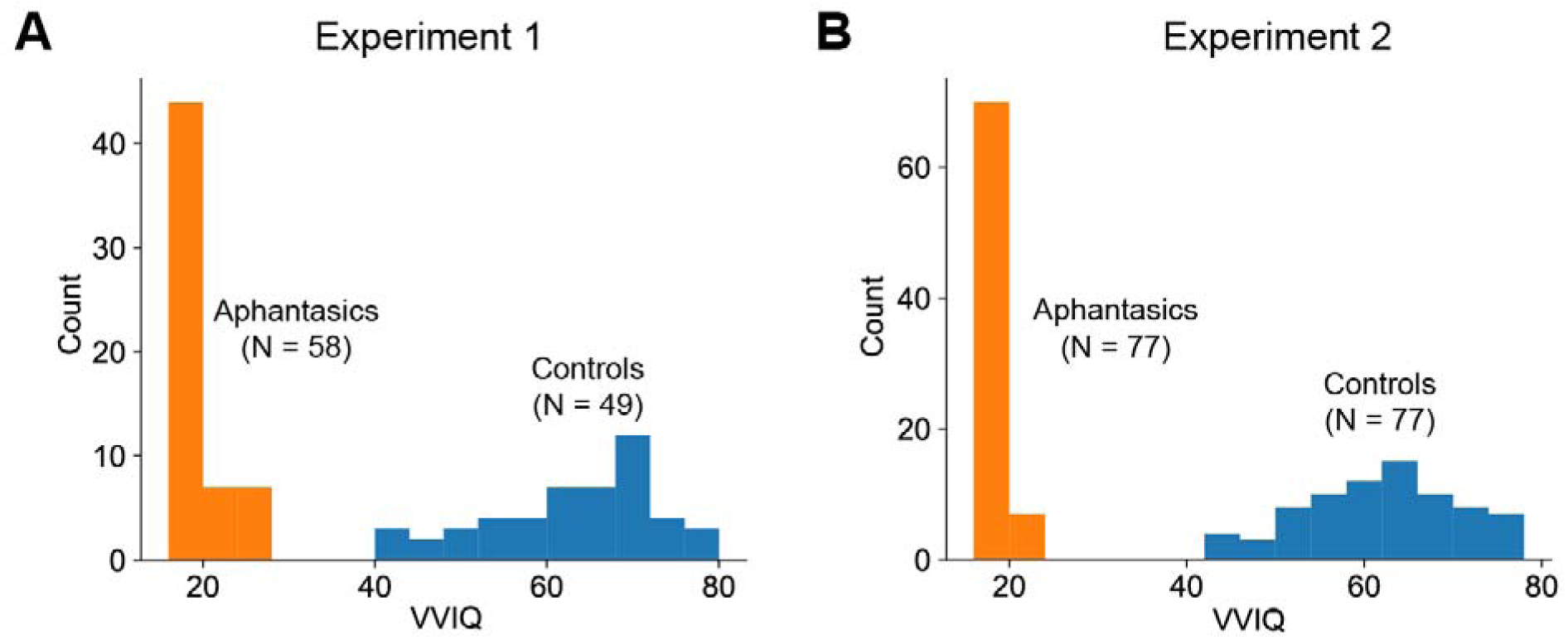
VVIQ of aphantasic (orange) and control (blue) subjects in Experiments 1 and 2. The two panels show the histograms of VVIQ distribution in Experiments 1 and 2, with aphantasic participants scoring 25 or below and control participants scoring 40 or above on the VVIQ scale.

Participants were presented with a total of five such attention checks throughout the task. To ensure data quality, only participants who answered all five questions correctly were retained for further analysis. In Experiment 1, we recruited 58 young adults with aphantasia (ages 18–35) from online communities such as the Aphantasia Network and relevant Reddit forums. Their average VVIQ score was 18.05 and a standard deviation of 3.18. The control group consisted of 49 participants recruited via Prolific, with an average VVIQ score of 62.47 and a standard deviation of 9.94.

#### Stimuli

For the color visual working memory task (**Fig 2**, upper panel), stimuli consisted of colored squares created using JavaScript and the jsPsych canvas-keyboard interface. Each square was 40 by 40 pixels and was displayed on a 800 by 800 pixel canvas.

**Fig 2.**
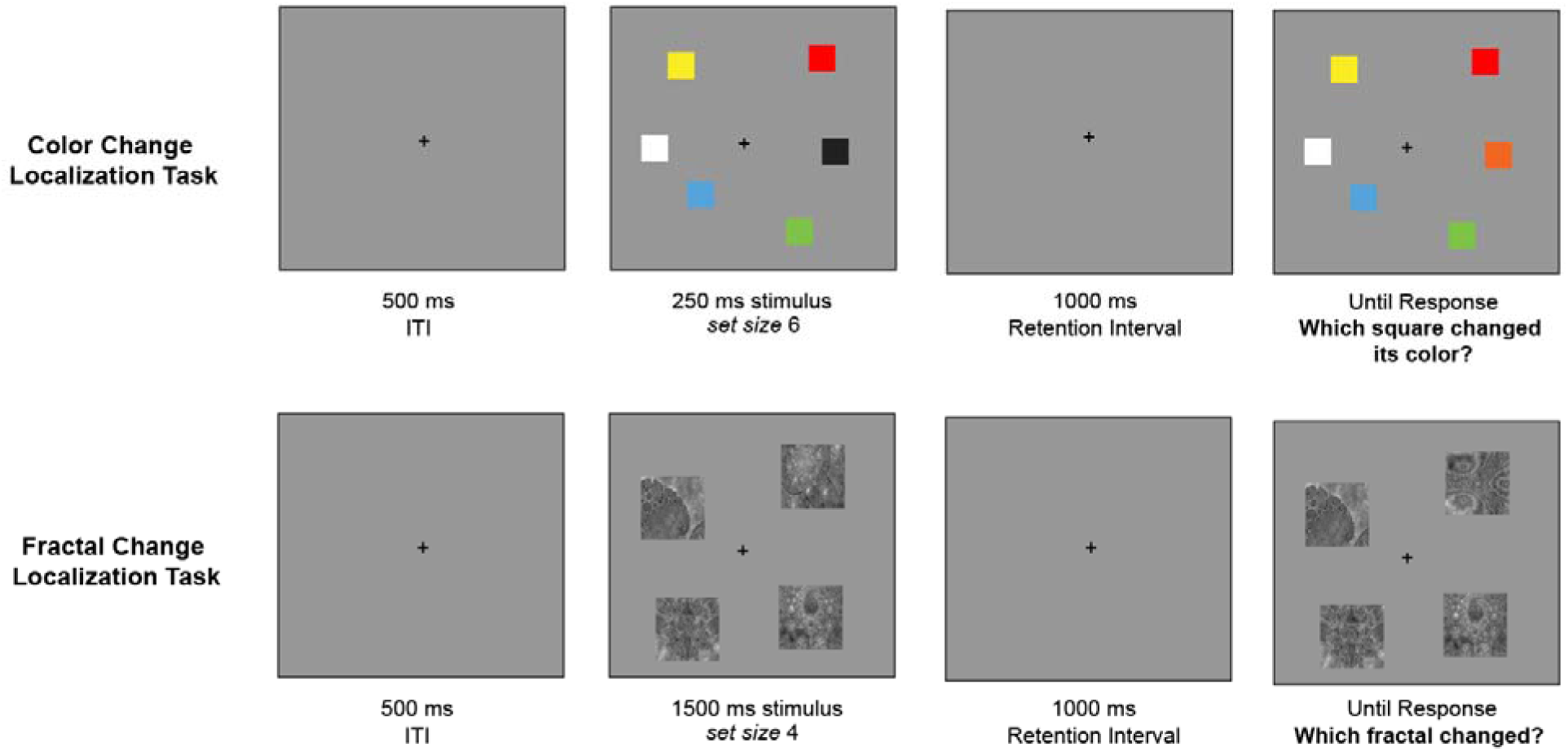
Color and Fractal Change Localization Paradigm used in Experiment 1. (Upper panel) Example trial from the color change localization task. Six colored squares are displayed briefly, followed by a delay, then a test display where one color has changed. (Lower panel) Example trial from the fractal change localization task. Four fractals are displayed for 1500 ms, followed by a delay, then a test display where one fractal has changed.

Squares appeared randomly within a circular region on the screen, positioned between 30 and 400 pixels from the canvas center. Each square was assigned one of nine unique colors per trial, with no repeats (RGB values: red = 255 0 0; green = 0 255 0; blue = 0 0 255; magenta = 255 0 255; yellow = 255 255 0; cyan = 0 255 255; orange = 255 128 0; white = 255 255 255; black = 0 0 0). For the fractal visual working memory task (**Fig 2**, lower panel), stimuli consisted of fractals selected from a standard fractal dataset (Ovalle-Fresa et al., 2022). To ensure that the number of possible fractals matched the number of color square options, we selected nine unique fractals from the full dataset and sampled four for each trial. In this dataset, the authors also collected nameability ratings for each image, indicating how likely participants were to assign a name to a given fractal. Each fractal was 60 by 60 pixels and was displayed on an 800 by 800 pixel canvas. Fractals appeared randomly within a circular region on the screen, positioned between 30 and 400 pixels from the canvas center. Throughout both tasks, participants were instructed to maintain fixation on a small black plus sign (30 px, Arial font) centered on the screen.

#### Procedure

For both the color and fractal visual working memory tasks, we used a change localization paradigm that had been shown to have high reliability and sensitivity with colors and shapes (Zhao et al., 2022). For the color change localization task, on each trial, six colored squares were shown at the same time for 250 milliseconds, followed by a 1,000-millisecond blank screen where participants had to retain the information in working memory. After this delay, the same six squares reappeared in their original positions, but one square had changed to a new color that had not been part of the initial display. Each square was labeled with a digit from 1 to 6, and participants identified the changed item by pressing the corresponding number key. In total, each participant completed 60 trials of the color change localization task (see **Fig. 2**, Upper panel).

For the fractal change localization task, on each trial, four fractals were shown at the same time for 1500 milliseconds, followed by a 1,000-millisecond blank screen where participants had to retain the information in working memory. For each trial, the four selected fractals were randomly selected from a set of nine preselected options drawn from the full dataset. Notably, we increased the presentation time and decreased the set size of our fractal stimuli compared to color squares used in the first block. The main reason here was that we would like most participants to perform above chance for the task to capture an accurate measure of capacity, and a pilot run with four individuals in our lab suggested that more than 75% of the participants showed below chance performance with six fractals presented for 250 ms. After the 1000-millisecond delay, the same four fractals reappeared in their original positions, but one fractal had changed to a new fractal that had not been part of the initial display. Each fractal was labeled with a digit from 1 to 4, and participants identified the changed item by pressing the corresponding number key. In total, each participant completed 60 trials of the fractal change localization task (see **Fig. 2**, Lower panel).

## Results

Our primary question here was to examine if aphantasics were impaired in visual working memory for color and fractal stimuli, where color stimuli are more nameable than the fractal stimuli. We found that aphantasics performed significantly worse than controls on the color visual working memory task (t(105) = -5.79, p < 0.01). Similarly, they showed impaired performance on the fractal working memory task (t(105) = -4.00, p < 0.01). These results suggest that aphantasic participants are worse in both forms of visual working memory tasks. To further examine whether the degree of impairment differed by task type (color vs. fractal), we conducted a mixed-effects linear model with Aphantasia and Task Type as predictors (accuracy ∼ aphantasia * task type).

Consistent with the t-test results, we observed a significant main effect of Aphantasia (β = -0.13, SE = .02, t(105) = -5.51, p < .01, **Fig 3**), indicating that aphantasics performed worse overall across visual working memory tasks. We also found a significant main effect of Task Type (β = .06, SE = .02, t(105) = 2.49, p = .01, **Fig 3**), with all participants performing worse on the fractal task compared to the color change localization task.

**Fig 3.**
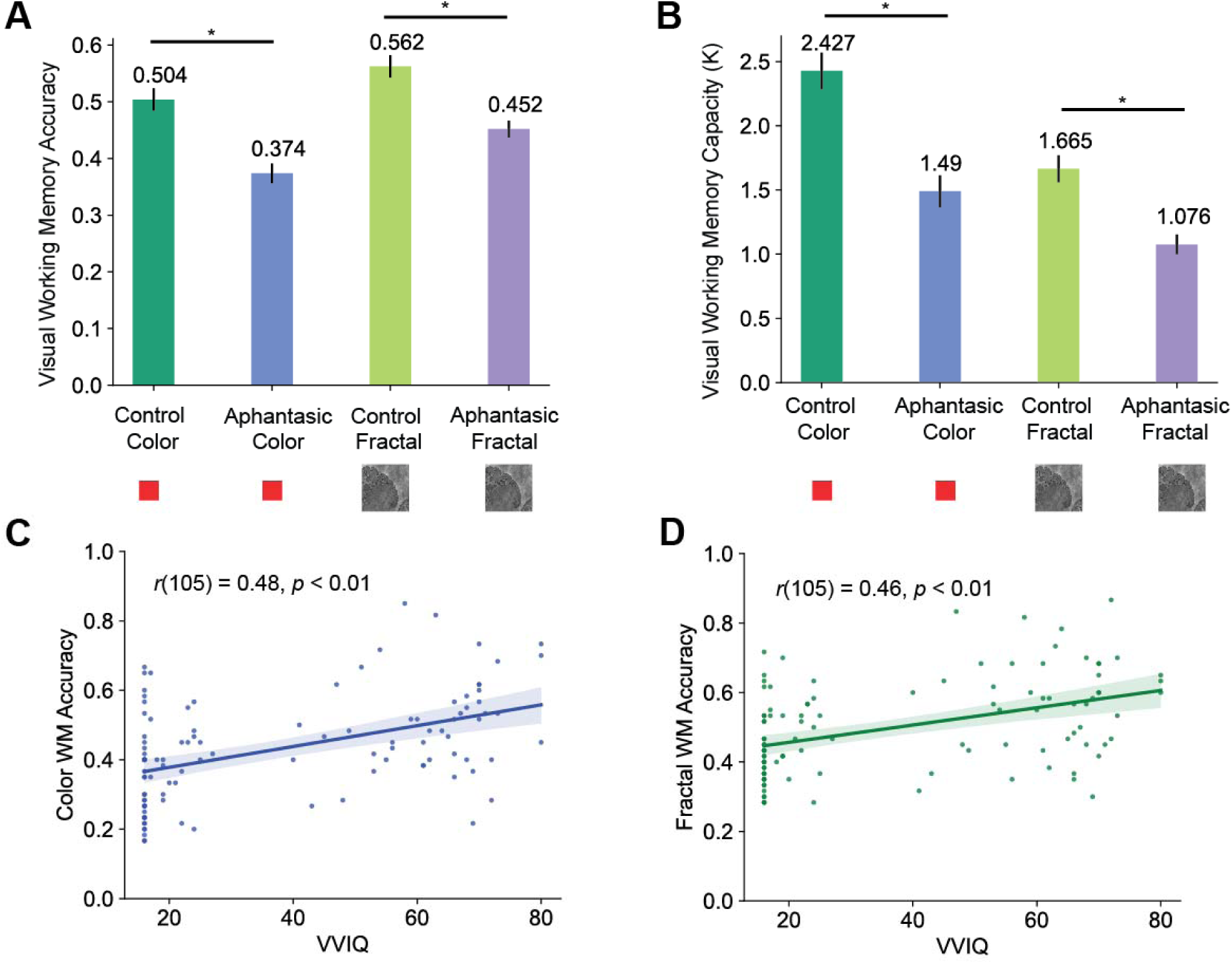
Working memory performance (A and B) and its relationship with VVIQ (C, D, and E) in Experiment 1. Aphantasic participants showed reduced working memory performance compared to controls on both the color and fractal tasks, as measured by accuracy (Panel A) and capacity (Panel B). Furthermore, individual differences in mental imagery ability, assessed by VVIQ scores, predicted performance on both tasks (Panels C and D). The error bars and shades reflected the standard error of the mean (SEM) of our data.

However, there was no significant interaction between Aphantasia and Task Type (β = .02, SE = .03, t(105) = 0.62, p = .54), suggesting that the impairment in aphantasics was not specific to one task type over the other. Within the fractal condition, we found that the average or maximum nameability of the fractal stimuli did not predict either control or aphantasic subjects’ memory performance (*p*s > 0.60), suggesting that the nameability of certain fractals did not affect working memory performance in either group. Collectively, these findings indicate that aphantasics have generally impaired visual working memory, regardless of stimulus type and regardless of whether they are more easily verbalizable (color) or less verbalizable (fractal) stimuli.

In addition to the accuracy of localization performance, researchers have proposed using a K capacity model, which estimates the number of items stored in visual working memory based on both hit and false alarm rates in change detection tasks (Cowan, 2001). Prior research has extended this formula to change localization tasks, where every trial involves a change, and demonstrated that the resulting K values closely match those obtained from traditional change detection paradigms (Zhao et al., 2022). Using this approach, we converted accuracy scores into K values to estimate visual working memory capacity

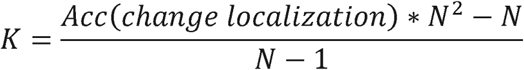

where N is set size. We then conducted a mixed-effects linear model with Aphantasia and Task Type as predictors (K ∼ aphantasia * task type). In line with our previous accuracy-based analyses, we observed a significant main effect of Aphantasia (β = -0.90, SE = .15, t(105) = -6.17, p < .01), indicating that aphantasics had reduced working memory capacity compared to controls. There was also a significant main effect of Task Type, with all participants performing worse on the fractal task than on the color change localization task (β = -0.76, SE = .15, t(105) = -5.01, p < .01). Importantly, we did not observe a significant interaction between Aphantasia and Task Type (β = .35, SE = .21, t(105) = 1.70, p = .09), suggesting that the deficit in aphantasics was not limited to a specific stimulus type. Taken together, both accuracy and K capacity measures support the conclusion that individuals with aphantasia exhibit a general impairment in visual working memory, regardless of the type of visual stimulus.

Lastly, we investigated whether participants’ mental imagery abilities could predict their visual working memory performance. To test this, we conducted correlational analyses between VVIQ scores, a measure of mental imagery vividness, and accuracy in both the color and fractal change localization tasks. Results showed that higher VVIQ scores were positively associated with greater accuracy in both color (*r*(105) = 0.48, p < .01, **Fig 3**) and fractal (*r*(105) = 0.46, p < .01, **Fig 3**) change localization tasks. A Fisher’s Z test comparing the two correlations indicated no significant difference between them (p = 0.9). Furthermore, we found that the control group VVIQ scores did not predict visual working memory performance with color squares (r(105) = 0.27, p = 0.06) or fractals (r(105) = 0.15, p = 0.31), suggesting that the correlations between mental imagery abilities and VWM were mainly driven by the group differences between aphantasics and controls. These findings suggest that individuals with stronger mental imagery abilities tend to perform better on visual working memory tasks, regardless of the type of stimuli.

## Discussion

In Experiment 1, our findings show that individuals with aphantasia are significantly impaired in visual working memory tasks involving both color and fractal stimuli. This impairment was evident across both accuracy and K capacity measures and did not differ by stimulus type, suggesting a general deficit in visual working memory rather than one tied to specific visual features. Additionally, we found that mental imagery ability, as measured by VVIQ, was positively correlated with performance in both tasks, supporting the idea that visual imagery contributes to visual working memory capacity.

While these results highlight a robust relationship between imagery and visual working memory, it remains unclear whether this deficit in aphantasia extends to non-visual domains. To address this question, Experiment 2 investigates whether the observed impairments also apply to visual working memory by using verbal stimuli such as words and pseudowords. A secondary question is whether aphantasic participants are able to use phonological coding to reduce their impairment due to the lack of visual imagery. To test this in Experiment 2, we use words, phonologically valid pseudowords, and phonologically invalid pseudowords as stimuli in a visual working memory task. If aphantasic participants rely on phonological coding to compensate for their impaired visual working memory, we would expect their performance to decline less in the word condition, which allows for semantic and phonological strategies, and the phonologically valid pseudoword condition, which supports phonological strategies, compared to the phonologically invalid pseudoword condition, which offers neither semantic nor phonological support.

## Experiment 2

### Methods

#### Participants

We used the same criteria for screening aphantasics and controls, and the same attentional check procedures, as in Experiment 1. In Experiment 2, we recruited 77 young adults with aphantasia (ages 18–35) from online communities such as the Aphantasia Network and relevant Reddit forums. Their average VVIQ score was 16.84 and a standard deviation of 1.90 (**Fig. 1B**). The control group consisted of 77 participants recruited via Prolific, with an average VVIQ score of 61.48 and a standard deviation of 8.80 (**Fig. 1B**).

#### Stimuli

The words and phonologically valid pseudowords used in Experiment 2 were selected from a database with words and phonologically valid pseudowords (Perrachione et al., 2017); see **Fig. 4** for examples. For the phonologically invalid pseudowords, to match the length of the stimuli to that of the other conditions, we randomly permuted letters of the word and phonologically valid pseudoword stimuli to produce a list of phonologically invalid pseudowords. We visually inspected the list such that none of the permutated stimuli were phonologically valid. The letters were presented with 40 pixel Arial font on an 800 by 800 pixel canvas. Words or pseudowords appeared randomly within a circular region on the screen, positioned between 50 and 400 pixels from the canvas center. Throughout all three tasks, participants were instructed to maintain fixation on a small black plus sign (30 px, Arial font) centered on the screen.

**Fig 4.**
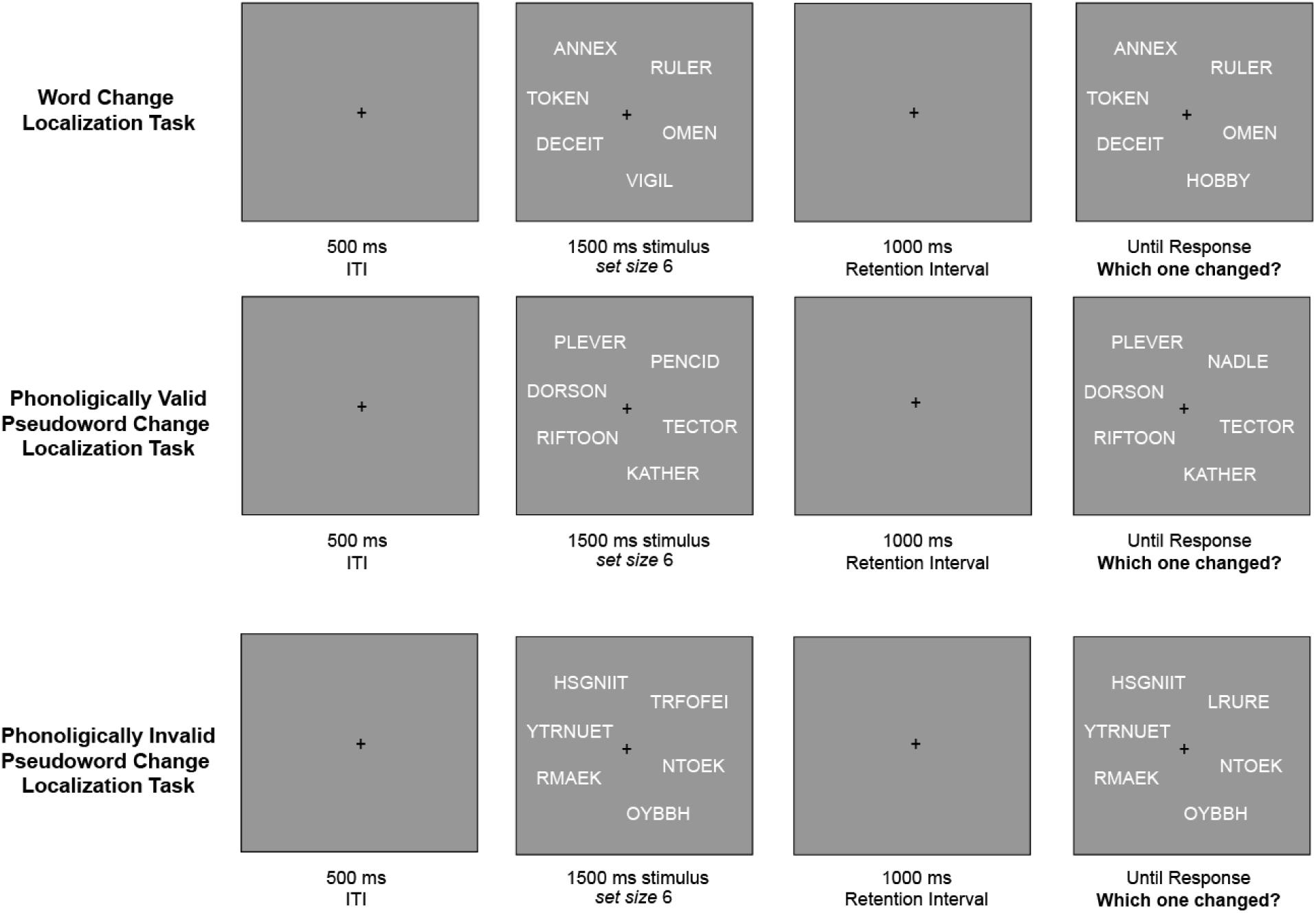
Word, Phonologically Valid Pseudoword and Phonologically Valid Pseudoword Change Localization Paradigm used in Experiment 2. (Upper panel) Example trial from the word change localization task. (Middle panel) Example trial from the phonologically valid pseudoword change localization task. (Lower panel) Example trial from the phonologically invalid pseudoword change localization task.

#### Procedure

For all three visual working memory tasks with verbal materials (words, phonologically valid pseudowords, and phonologically invalid pseudowords), we used the change localization paradigm used in Experiment 1. On each trial, six words, phonologically valid pseudowords or phonologically invalid pseudowords were shown at the same time for 1500 milliseconds, followed by a 1,000-millisecond blank screen to retain the information. After this delay, the same six words, phonologically valid pseudowords or phonologically invalid pseudowords reappeared in their original positions, but one stimulus had changed to a new word, phonologically valid pseudoword or phonologically invalid pseudoword that had not been part of the initial display. Each square was labeled with a digit from 1 to 6, and participants identified the changed item by pressing the corresponding number key. In total, each participant completed 60 trials of each type of change localization task (see **Fig. 4**).

## Results

In Experiment 2, our primary goal was to generalize our findings in Experiment 1 to words and non-word letter strings with three conditions: the word condition (testing use of phonological and semantic strategies), the phonologically valid pseudoword condition (testing use of phonological strategies), and the phonologically invalid pseudoword condition (providing minimal semantic or phonological support). To test this, we conducted an independent-samples t-test on the accuracies for all three tasks between aphantasics and control subjects. We found that aphantasics performed significantly worse than controls on the word change localization task (t(152) = -6.40, p < 0.01, **Fig 5**). Similarly, they showed impaired performance on the change localization tasks with phonologically valid pseudowords (t(152) = -4.84, p < 0.01, **Fig 5**) and phonologically invalid pseudowords (t(152) = -4.10, p < 0.01, **Fig 5**). To examine whether the degree of impairment differed by task type (words, phonologically valid pseudowords, and phonologically invalid pseudowords), we conducted a mixed-effects linear model with Aphantasia and Task Type as predictors (accuracy ∼ aphantasia * task type). Consistent with the t-test results, we observed a significant main effect of Aphantasia (β = -0.13, SE = .03, t(152) = -4.22, p < .01), indicating that aphantasics performed worse overall across visual working memory tasks. We also found a significant main effect of Task Type (β = .06, SE = .02, t(152) = 4.28, p = .01), with all participants performing worse on the phonologically invalid pseudoword change localization tasks compared to the word task (t(152)s > 4.23, ps < .01). Moreover, the aphantasic participants performed worse on the phonologically invalid pseudoword change localization task than phonologically valid pseudoword task (t(152) = 2.40, p = 0.02), while the control subjects performed similarly in these two tasks (t(152) = 0.98, p = 0.33). However, there was no significant interaction between Aphantasia and Task Type (β = .03, SE = .02, t(152) = 1.64, p = .10), suggesting that the impairment in aphantasics was not specific to one task type over the other. Collectively, these findings indicate that aphantasics have generally impaired visual working memory for verbal materials, regardless of whether the materials are meaningful (words versus pseudowords) or phonologically valid (words and phonologically valid pseudowords versus phonologically invalid pseudowords).

**Fig 5.**
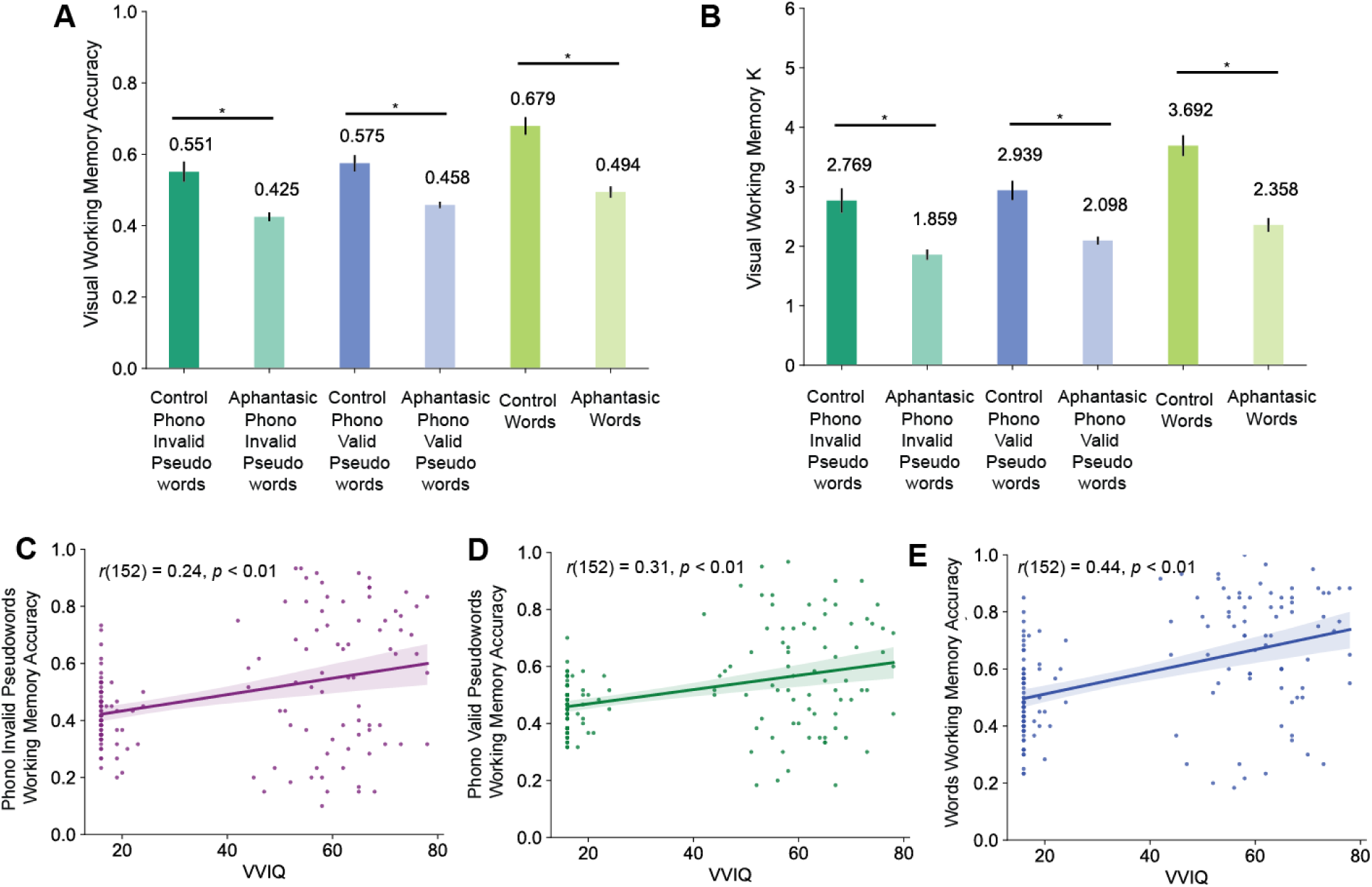
Working memory performance (A and B) and its relationship with VVIQ (C, D, and E) in Experiment 2. Aphantasic participants showed reduced working memory performance compared to controls on all three tasks, as measured by accuracy (Panel A) and capacity (Panel B). Furthermore, individual differences in mental imagery ability, assessed by VVIQ scores, predicted performance on all three tasks (Panels C, D, and E). The error bars and shades reflected the standard error of the mean (SEM) of our data.

Lastly, we investigated whether participants’ mental imagery abilities could predict their visual working memory performance for verbal materials. To test this, we conducted correlational analyses between VVIQ scores, a measure of mental imagery vividness, and accuracy in all three versions change localization tasks. Results showed that higher VVIQ scores were positively associated with greater accuracy in word (r(152) = 0.44, p < .01, **Fig 5**), phonologically valid pseudoword (r(152) = 0.31, p < .01, **Fig 5**), and phonologically invalid pseudoword (r(152) = 0.24, p < .01, **Fig 5**) change localization tasks. Interestingly, Fisher’s Z tests comparing the correlation strength suggested that VVIQ predicted word change localization performance better than phonologically invalid word performance (z(152) = 2.32, p = .01). Similar to Experiment 1, we found that VVIQ scores for the control group did not predict their visual working memory performance for words (r(152) = -0.02, p = 0.90), phonologically valid pseudowords (r(152) = 0.03, p = 0.76), or phonologically invalid pseudowords (r(152) = 0.12, p = 0.29), suggesting that the correlations between mental imagery abilities and VWM were mainly driven by the group differences between aphantasics and controls. Our findings suggested that mental imagery ability was a better predictor for visual working memory with meaningful and verbalizable materials than with meaningless and hardly verbalizable materials. Collectively, our findings extended Experiment 1’s findings into visual working memory tasks using verbal materials, and showed that individuals with stronger mental imagery abilities tend to perform better on visual working memory tasks even with verbal materials.

## Discussion

Experiment 2 aimed to investigate whether the visual working memory impairment observed in aphantasics for non-verbal stimuli would also extend to verbal materials. Consistent with our findings in Experiment 1, aphantasics performed significantly worse than control participants across all verbal visual working memory tasks, including words, phonologically valid pseudowords, and phonologically invalid pseudowords. This impairment was consistent across conditions, with no significant interaction between Aphantasia and Task Type, indicating that the deficit was not modulated by the phonological properties or meaningfulness of the stimuli.

Crucially, these results suggest that aphantasics’ impairment in visual working memory persists even when stimuli are verbal or phonologically codable. If aphantasics were compensating for their imagery deficits using phonological strategies, we would have expected a relative performance benefit for phonologically valid or familiar items. However, no such effect was observed, implying that phonological coding does not substantially reduce the visual working memory impairment in aphantasics.

Moreover, mental imagery ability, as measured by VVIQ scores, continued to predict performance in these tasks, with stronger correlations for more meaningful and verbalizable stimuli. These findings extend the results of Experiment 1 and further support the view that mental imagery plays a functional role in visual working memory even for verbal materials. Overall, our data suggest that individuals with aphantasia experience a broad impairment in visual working memory that is not easily compensated for by alternative strategies such as phonological encoding.

## General Discussion

The present study investigated the relationship between visual imagery and visual working memory (VWM), two cognitive functions that are often conceptually intertwined. Across two experiments, we examined VWM performance in aphantasic and control participants using a change localization task with a range of stimulus types. In Experiment 1, we tested memory for color squares and complex fractals; in Experiment 2, we extended this to include another category of visual stimuli: real words, phonologically valid pseudowords, and phonologically invalid pseudowords. Aphantasic participants consistently showed impaired VWM performance compared to controls, regardless of the stimulus type. Importantly, their deficits were evident for both easily verbalizable stimuli (e.g., colors, words) and those that are difficult to verbalize (e.g., fractals, pseudowords), suggesting that compensatory verbal strategies were insufficient to support performance. Collectively, our experiments indicated that aphantasic participants had impaired visual working memory capacity across a wide range of stimuli.

Our study suggested that visual imagery plays a significant role in visual working memory capacity. Our behavioral findings support previous neuroscience research indicating that these two processes may rely on similar brain regions. For example, a prior study asked participants to either encode orientation patterns into visual working memory or to imagine the same patterns while nothing was shown on the screen.

Activity patterns in early visual areas (V1–V3) allowed for successful decoding of the orientation under both the working memory and imagery conditions (Albers et al., 2013). The overlap between these processes suggests that individuals with lower visual imagery abilities may show different activation patterns in these regions. A more recent study compared the neural patterns of an aphantasic individual and their control twin during a visual imagery task (Megla et al., 2025). Multivariate analyses revealed that the aphantasic twin showed reduced connectivity between occipital and temporal lobes, suggesting that early visual areas may support the representation of visual information in both memory and imagery. Future research involving a larger sample of aphantasic participants is needed to identify which brain regions are responsible for impaired visual imagery, and which patterns of connectivity may explain deficits in visual working memory related to the absence of imagery.

One important finding from our experiment is that visual working memory appears to store visually presented materials including color squares, fractals, words and non-word letter strings, and were similarly impaired in people with lower imagery abilities. This aligns with prior neuroscience research that examined visual working memory using verbal stimuli. In typical visual working memory tasks involving visual items such as colored squares, contralateral delay activity (CDA), a sustained negativity observed over occipital-parietal regions contralateral to the side of the stimuli, increases in amplitude as more items are held in memory (Vogel & Machizawa, 2004). Notably, CDA amplitude plateaus once an individual’s visual working memory capacity is reached, providing a reliable neural index of memory load. A recent study extended this work by testing whether letters and words, which are typically associated with phonological processing, would elicit similar CDA patterns (Rajsic et al., 2019). They found that both letters and words evoked a load-sensitive CDA, suggesting that these verbal stimuli are encoded visually in working memory, rather than relying purely on phonological representations. Like with non-verbal stimuli, the CDA amplitude increased with memory load and leveled off at capacity, indicating similar underlying mechanisms. Our study adds to this body of work by showing that visual working memory performance was impaired in individuals with aphantasia, regardless of whether the materials were visual or verbal. This provides compelling behavioral evidence that supports the theory that visual encoding plays a central role in visual working memory, across different types of stimuli.

An open question not addressed by our study is whether the absence of visual imagery may lead to broader deficits in general cognitive abilities. One major area of interest is visual memory. Prior research has shown that aphantasics perform worse on object memory tasks, but not on spatial memory tasks, when asked to recall drawings from long-term memory (Bainbridge et al., 2021). Given the well-established relationship between working memory and long-term memory (Adam et al., 2024; Fukuda & Vogel, 2019; Robison et al., 2024; Unsworth et al., 2014; Zhao & Vogel, 2025b, 2025a), future research is needed to determine whether the observed object memory deficits in aphantasics stem from impairments in visual working memory capacity. In other words, if aphantasics encode less visual information into working memory initially, they may have fewer representations available for consolidation into long-term memory.

Another compelling question is whether aphantasics also show deficits in sustained visual attention as a result of impaired visual working memory. For example, in tasks such as the gradual Continuous Performance Task (gradCPT, M. Rosenberg et al., 2013; M. D. Rosenberg et al., 2020), participants respond to a continuous stream of visual stimuli (e.g., images of indoor and outdoor scenes) by pressing a button for frequent stimuli (e.g., outdoor scenes shown 90% of the time) and withholding their response for infrequent ones. If visual working memory is compromised, we might expect diminished performance on such sustained attention tasks. Supporting this idea, recent findings suggest that long-term memory contributes to sustained attention in control participants (Zhao et al., 2025). Given the established deficits in long-term object memory among aphantasics (Bainbridge et al., 2021), it is worth investigating whether their sustained attention is also disproportionately affected due to limitations in visual long-term memory.

Together, our study explored the relationship between visual imagery and visual working memory (VWM) across two experiments using a variety of stimulus types.

Aphantasic individuals, who report little to no visual imagery, consistently showed reduced VWM performance compared to control participants. Notably, these deficits were not limited to a specific type of material; they generalized across both easily verbalizable stimuli (such as colors and real words) and more difficult-to-verbalize stimuli (such as complex fractals and pseudowords). This suggests that simple verbal strategies were insufficient for aphantasic individuals to compensate for their lower VWM performance. Furthermore, across both experiments, subjective imagery strength as measured by the VVIQ (Vividness of Visual Imagery Questionnaire) was positively correlated with VWM performance for all types of stimuli. This finding points to a potential general role of visual imagery in supporting visual working memory, beyond just tasks involving picturable or nameable content. Together, these results provide converging evidence that visual imagery may play a broad and functional role in visual working memory. Our findings highlight the importance of considering individual differences in mental imagery when investigating the mechanisms underlying visual working memory.

## Acknowledgements

This research was supported by the funding from National Institute of Mental Health (grant ROIMH087214 to E. K. V.); Office of Naval Research (grant N00014-12-1-0972 to E. K. V.); National Eye Institute (R01-EY034432 to W.A.B.).

